# Synaptic signaling networks encode experience by assuming stimulus-specific and brain-region-specific states

**DOI:** 10.1101/2021.03.25.437050

**Authors:** Jonathan D. Lautz, Kaleb B. Tsegay, Zhiyi Zhu, Edward P. Gniffke, John P. Welsh, Stephen E.P. Smith

## Abstract

A core network of ubiquitously expressed glutamate-synapse-associated proteins mediates activity-dependent synaptic plasticity throughout the brain, but the specific proteomic composition of synapses differs between brain regions. Here, we sought to classify the diversity of activity-dependent remodeling across brain regions using quantitative protein interaction network (PIN) analysis. We first compared the response of cultured neurons to distinct stimuli, and defined PIN parameters that differentiate input types. We next compared the response of three different brain regions maintained alive in vitro to an identical stimulus, and identified three qualitatively different PIN responses. Finally, we measured the PIN response following associative learning tasks, delay and trace eyeblink conditioning, in three brain regions, and found that the two forms of associative learning are distinguished from each other using brain-region-specific network mechanisms. We conclude that although the PIN of the glutamatergic post-synapse is expressed ubiquitously, its activity-dependent dynamics show remarkable stimulus-specific and brain-region-specific diversity.

## Introduction

Proteomic characterization of the glutamatergic postsynapse has identified thousands of ‘synaptic’ proteins in brain-wide preparations [1–6], as well as considerable variation between brain regions [7–9]. While core synaptic scaffolds (e.g. PSD95, Homer, Shanks) or receptors (e.g. AMPAR or NMDAR) are present at most excitatory synapses, hundreds of additional proteins are differentially expressed across brain regions [10–12]. These proteins assemble into large multiprotein complexes, sometimes mega-Dalton in size [13–16], the composition of which also shows considerable heterogeneity across brain regions, developmental stages, or downstream of genetic mutations [17, 18]. This protein-level diversity is believed to underlie morphological [19] and electrophysiological [20–22] diversity among neurons that receive glutamatergic input, and to be fundamental to the complexity and information processing capacity of the brain.

Downstream of synaptic activity, modifications to the proteome change the synapse’s response to subsequent activity in a process generally referred to as synaptic plasticity. Measured electrophysiologically, there is great diversity in plasticity, with different cell types responding in different ways to identical stimuli [23–25]. However, measured biochemically, the level of diversity is less clear. Activation of CamKII [26], mobilization of SynGAP [27–29] phosphorylation of NMDARs [30–32], and release of mGluR5 from Homer scaffolds [29, 33, 34] have been described for hippocampal or neocortical neurons, and likely occur elsewhere. At a proteomic level, induction of long-term potentiation (LTP) in the hippocampus has been shown to alter the phosphorylation status of 570 sites across 220 postsynaptic proteins [35]. Moreover, pharmacological manipulation of ionotropic, metabotropic, and dopamine receptors in the hippocampus activates overlapping phosphorylation networks, such that distinct physiological inputs activate different combinations of biochemical pathways [35]. Similarly, in neocortical neurons, stimulation of ionotropic vs. metabotropic glutamate receptors elicits distinct changes in synaptic protein complexes [29]. Collectively, these data support the hypothesis that different combinations of proteins in complex with each other encode units of information, such that discrete signaling inputs should trigger different combinatorial protein-state-codes, originally proposed by Pawson and colleagues in 2000 [36]. However, weather these codes remain consistent across different brain regions is not known.

Here, we address the question, how generalizable is the protein-state-coding of experience in the brain? Are glutamatergic postsynapses with distinct proteomic compositions and/or organizations modified by a similar set of ‘molecular logic’ rules? Or alternatively, do differentially expressed proteins fundamentally change the rules of molecular logic circuits, such that identical inputs lead to fundamentally different responses? We characterize the state of a glutamatergic synapse protein interaction network (PIN) following discrete molecular inputs, including acute stimulation of glutamate receptors and chemical induction of synaptic plasticity in vitro, and the induction of two different forms of associative learning measured in three brain areas of the mouse following classical eye blink conditioning (EBC). We demonstrate that discrete signaling inputs induce unique PIN states, and that responses to the same stimulus are often quite different across brain areas due to differences in the composition of the protein network and the behavioral context in which the stimuli are received. We posit that these combinatorial PIN states underlie the high-information processing capabilities of the synapse and that the diversity of PIN responses enables different brain regions to respond to and integrate a broad range of signaling inputs.

## Results

### NMDA, DHPG and Glutamate stimulation produce widespread PIN re-arrangement

PIN networks can be conceptualized as molecular logic circuits, simultaneously translating multiple receptor inputs into intracellular signals and performing molecular calculations to integrate inputs and select an appropriate response (Fig 1A). In previous studies, we developed a system to investigate the molecular logic coding of a PIN composed of glutamate receptors, scaffolds, and downstream signaling proteins, called quantitative multiplex co-immunoprecipitation (QMI) [29]. QMI uses antibody-coupled flow cytometry beads to quantify changes in targeted Proteins in Shared Complexes with Exposed Epitopes (PiSCES); we refer to detected interactions as PiSCES instead of protein-protein interactions to highlight that the detected interactions are not necessarily direct. Our QMI panel is composed of 20 protein targets selected based on synaptic expression, known co-association, and linkage to autism, and includes Glutamate receptors (NMDA subunits NR1, 2A, 2B, AMPA subunits GluR1 and GluR2, and metabotropic glutamate receptor (mGluR5)), scaffolding proteins (Shank 1 and 3, Homer 1 and 1a, PSD95, SAP97, SAPAP1, Neuroligin3), and signal transduction effectors (SynGAP, Fyn, CamKII, PI3K, PIKE, and Ube3A) [29, 37]. Importantly, QMI has been validated for antibody specificity [29] and detergent-independence [38].

**Figure 1:**
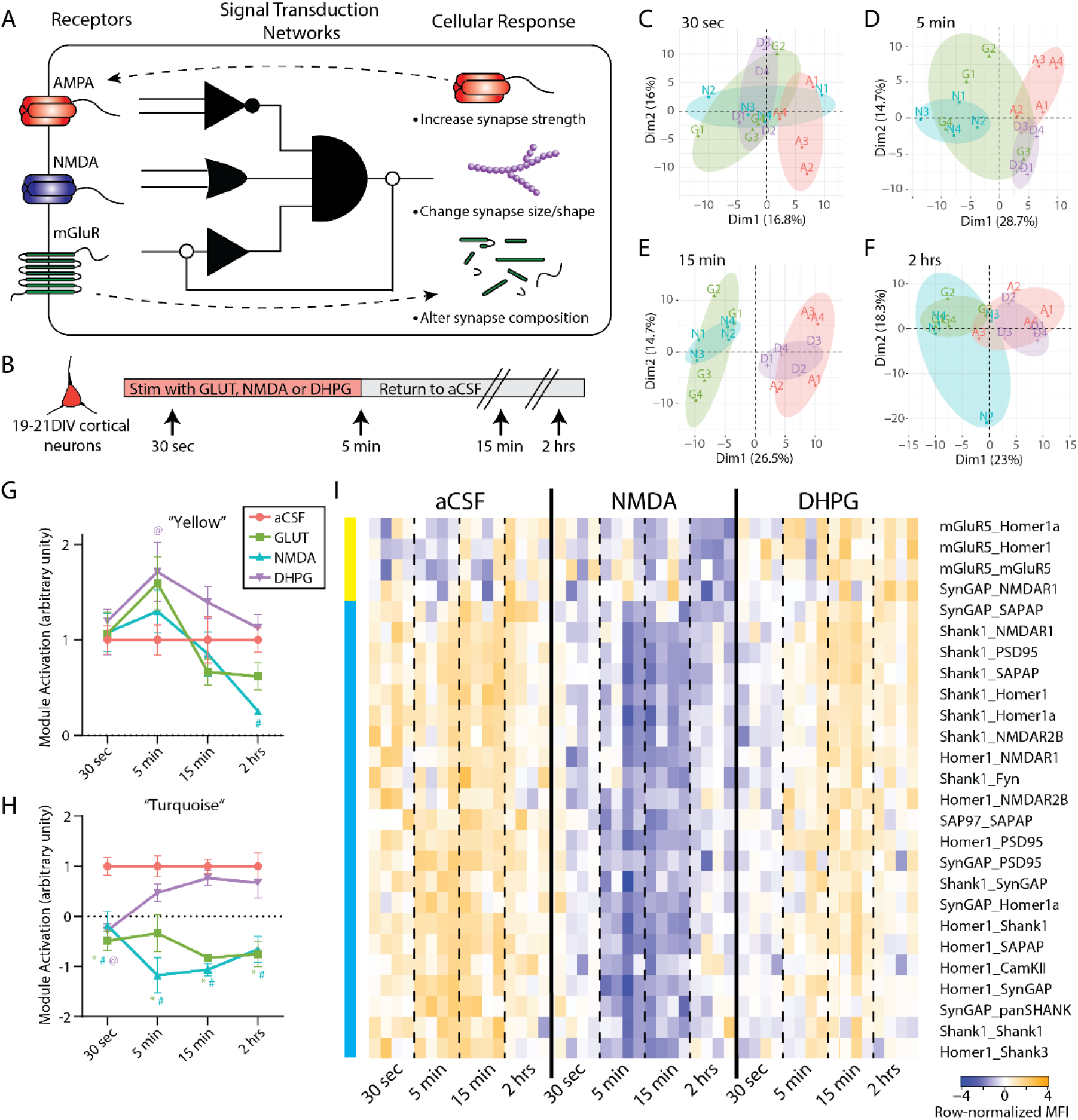
Timecourse of NMDA and DHPG stimulation. A) Conceptual illustration of a PIN logic circuit linking receptor inputs with cellular outcome. B) Experimental design. C-F) PCA of 30 (C) second timepoint, 5min (D), 15min (E) and 2 hour (F) timepoints shows separation of aCSF [A], glutamate [G], NMDA [N] and DHPG [D]. G) Average normalized MFI of PiSCES in the turquoise CNA module over the aCSF, Glutamate, NMDA and DHPG timecourse. N=4 per timepoint per treatment. *, # and @ indicate p<0.005 for Glut, NMDA and DHPG, respectively, by Dunnet’s multiple comparison following 2-way ANOVA (Treatment F_3,48_=52.31, p<0.0001, Time F_3,48_=0.13 NS, Interaction F_9,48_=2.92, p<0.01). H) Average normalized MFI of PiSCES in the yellow CNA module. N=4 per timepoint per treatment. *, # and @ indicate p<0.05 for Glut, NMDA and DHPG, respectively, by Dunnet’s multiple comparison following 2-way ANOVA (Treatment F_3,48_=4.78, p<0.01, Time F_3,48_=7.94, p<0.0005, Interaction F_9,48_=1.65, NS). I) Heatmap of all PiSCES ANC∩CNA significant the entire time course. The CNA modules are illustrated by colored bars on the left of the figure. Data are expressed as row-normalized MFI.

Prior data suggest that the glutamate synapse PIN uses a qualitative mechanism to encode distinct signaling inputs: different PiSCES are engaged by five minutes of NMDA vs. mGluR stimulation [29]. However, in other systems, quantitative (i.e. intensity) [39] or kinetic [40] differences in the engagement of *identical* signaling intermediates allow cells to distinguish between input types. To further characterize PIN logic-coding in a simple model system, we stimulated DIV19-21 cortical neuronal cultures for up to 5 minutes with glutamate (GLUT), NMDA/glycine, DHPG, or aCSF control and measured the PIN response at 30s, 5 min, 15 min, and 2 hr to establish the qualitative, quantitative and kinetic differences between different types of synaptic stimulation (Fig 1B).

To ensure cells remained healthy and active following treatment, we performed Ca2+ imaging of cultures for 30 min following stimulation. Treatment with glutamate or NMDA/glycine induced a rapid increase in intracellular Ca^2+^ (Fig. S1A,B), whereas we were unable to detect any significant changes in intracellular calcium staining following treatment with DHPG. Upon removal of agonists, intracellular Ca2+ returned to baseline for 30 min, and subsequent depolarization with KCL increased intracellular Ca^2+^, demonstrating that cells remained alive and receptive to stimuli (Fig S1A,B). We also used TUNEL staining to quantify agonist-induced apoptosis at 2h and did not observe increased rates neuronal apoptosis in any condition (Fig S1C,D).

To visualize high-dimensional QMI data, we first applied principal component analysis (PCA). After 30S s, all three treatment conditions separated from aCSF across PC1 (Fig 1C). At 5 min, glutamate and NMDA stimulation continued to separate from aCSF across PC1, while DHPG normalized across PC1 but separated across PC2 (Fig 1D), similar to our previous report on only the 5 min timepoint [29]. By 15min, aCSF and DHPG overlapped, while NMDA and glutamate remained separated across PC1 (Fig 1E). At 2 hr, the separation between aCSF/DHPG and NMDA/glutamate, while still present, was smaller in magnitude (Fig 1F). Overall, NMDA and glutamate stimulation produced a stronger, more sustained response, whereas DHPG induced a transient response that was initially similar to NMDA/GLUT but diverged at 5 min.

To further explore network-scale temporal dynamics, we performed weighted correlation network analysis (CNA) on all timepoints. Out of 193 pairwise PiSCES measurements above background comprising 6 CNA modules, two modules were significantly correlated with a treatment. One module (colored turquoise in Fig 1H, I) correlated with all treatments, and contained 22 PiSCES that met the strict criteria of module membership (MM) > 0.7. All turquoise PiSCES were also significant by ANC, a second statistical test that relies on independent assumptions [41]. To compare turquoise module behavior across stimulations, we averaged the z-scaled intensity of PiSCES for each treatment/timepoint (n = 4 each) and normalized to aCSF (Fig 1H). At 30 s, all three treatment conditions were significantly different from aCSF. In DHPG, the module was no longer significantly different by 5 min, but in NMDA and GLUT, the module activation continued to strengthen, peaking at 5-15 min, and remaining significantly different even after 2 hr. Thus, while the “turquoise module” was activated in all treatment conditions, the kinetics of activation distinguished NMDA/GLUT (strong and sustained) from DHPG (weak and transient). A second CNA module, ‘yellow’, was correlated with only DHPG treatment, and its z-scaled intensity was significantly different from aCSF in the DHPG condition, only at 5 min (Fig 1G), although GLUT stimulation (which targets both NMDA and mGluRs) trended towards significance as well.

We next looked more closely at the specific PiSCES contained within each module by row-normalizing the median fluorescent intensity (MFI) of each ANC∩CNA PiSCES in the Yellow or Turquoise modules and displaying them as a heatmap for aCSF, NMDA and DHPG treatment (Fig 1I). These data reveal a rapid dissociation of Homer, Shank, and SynGAP-containing PiSCES at 30 s, which increased in magnitude at 5 and 15 min in NMDA, but rapidly returned to baseline in DHPG. Even after 2 hr, PiSCES remained lower in NMDA and DHPG conditions. ANC∩CNA analysis was also performed on each timepoint independently (Fig S2), and revealed additional PiSCES important to synaptic plasticity, such as increased association among PSD95 and AMPA-type glutamate receptors (GluR1/2), and changes in NMDAR interactions. By 2 hr, elevated levels of several NMDAR1 interactions, as well as PSD95_PSD95 and Homer1_Homer1 was apparent, reflecting oligomerization or expansion of synaptic scaffolding networks.

Overall, these data highlight two PiSCES modules, centered on Homer-, Shank-, SynGAP-, and mGluR5 containing interactions, that activate following glutamate agonist stimulation. DHPG vs. NMDA stimulation is encoded within these modules by quantitative (yellow ON/OFF), qualitative (turquoise weak/strong), and temporal (turquoise fast/slow) parameters.

### Chemical LTP activates a different network of PiSCES

Long term potentiation (LTP) is perhaps the most well studied form of synaptic plasticity and is induced electrophysiologically by pulsed activations of excitatory synaptic inputs. Activating downstream signaling pathways directly with small molecules using ‘chemical LTP’ (cLTP) has been shown to mimic many of the electrophysiological and biochemical features of LTP in cultured neurons [42–44]. We applied a commonly used cLTP protocol [44] to DIV 19-21 cultured cortical neurons and performed QMI. PCA revealed separation of control and cLTP neurons (Fig 2A), and ANC∩CNA analysis identified 20 PiSCES that were significantly altered (Fig 2B), consisting of increased Shank1-, Homer1-, and SynGAP containing PiSCES but decreased NL3, PIKE, and Fyn PiSCES. We compared these data to the GLUT 15-min timepoint because it most closely matched the time of the cLTP treatment (30 min), and involved the activation of the greatest number PiSCES in the agonist experiments. When the log2 fold change of all ANC∩CNA PiSCES were plotted on an X-Y axis, cLTP and GLUT application produced strikingly different patterns of PiSCES activation (Fig 2C). Nine of 20 PiSCES activated by cLTP were not activated by GLUT, and 23/32 PiSCES activated by GLUT were not activated by cLTP. Of the 11 PiSCES activated by both treatments, the majority (8/11) changed in the opposite direction (e.g. Shank1_Homer1a was strongly decreased in GLUT but increased in cLTP, #1 in Fig 2C). These data demonstrate the synaptic PIN under study can adopt multiple input-specific states in a cell culture paradigm.

**Figure 2:**
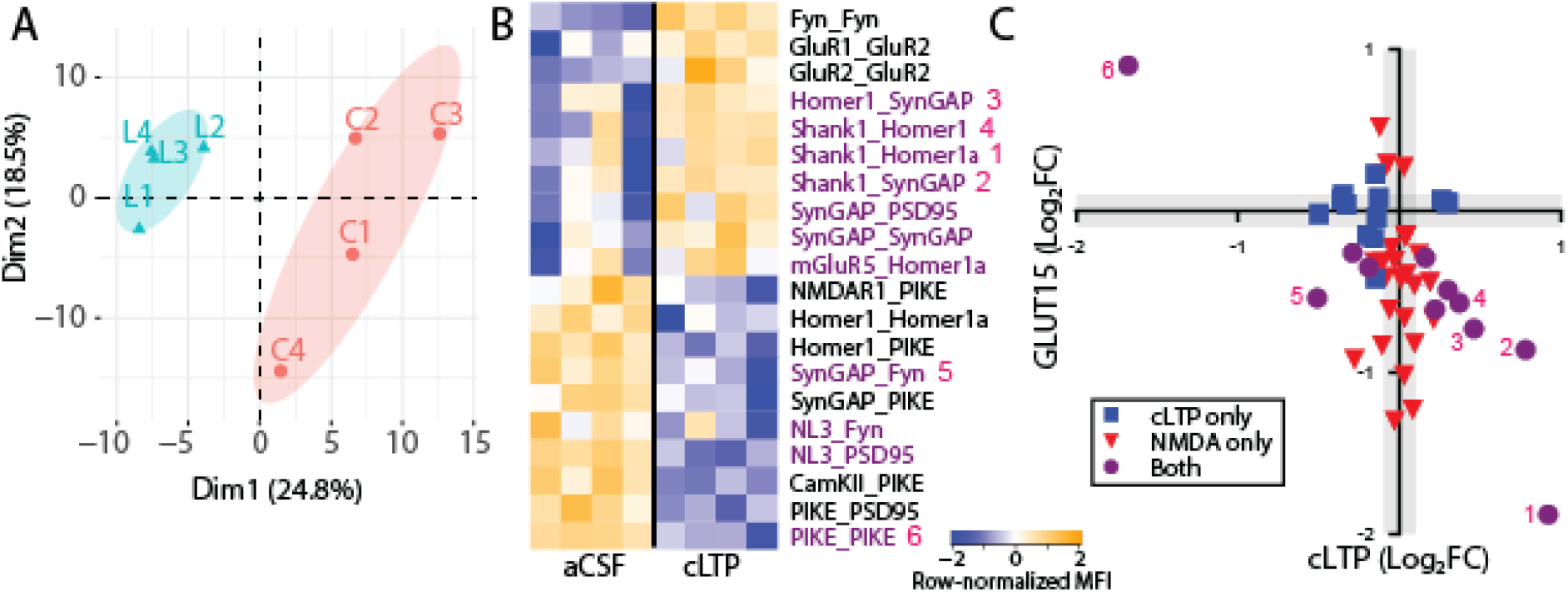
Chemical LTP. A) PCA shows control [C] and cLTP [L]. B) Heatmap of all PiSCES ANC∩CNA significant for the cLTP vs. control comparison. Data are expressed as row-normalized MFI. C) X-Y plot comparing the log_2_ fold change (FC) of all PiSCES that were ANC∩CNA significant for either the cLTP experiment (blue), the 15minute Glutamate timepoint (red) or both experiments (purple).

### NMDA stimulation of acute brain slices reveals differences across brain areas

We next asked how neurons with known differences in proteomic composition would respond to the *same* stimulus. We microdissected cortical (CTX) or hippocampal (HC) tissue from acute slice preparations, stimulated with NMDA/glycine or aCSF control for 5 min, and ran QMI in parallel to directly compare results in the two tissue types (Fig 3A). PCA of QMI data showed separation between CTX and HC, confirming that the proteomic organization of the two tissues is distinct (Fig 3B). Directly comparing only CTX vs. HC tissue (ignoring NMDA treatment) by ANC∩CNA, we identified 22 PiSCES that were significantly different between tissues (Fig 3C); notable differences included AMPA receptor PiSCES GluR1_GluR2 and GluR1_GluR1 (higher in hippocampus), AMPA-PSD95 associations (higher in cortex), multiple Homer-containing PiSCES (higher in hippocampus, although Homer self-association was higher in cortex), and PI3K/PIKE/NMDAR PiSCES (higher in hippocampus). However, the large PiSCES differences between tissue types masked the more subtle activity-dependent differences within each tissue. We therefore limited our analysis to comparisons of aCSF vs NMDA within the same tissue type using ANC∩CNA, and visualized the differences with heatmaps (Fig 3D). In CTX, 14 PiSCES changed significantly with NMDA, and 10 changed in HC (5 overlapping). Of these activity-dependent PiSCES, 8/19 were also significantly different in HC vs. CTX in the aCSF condition. Only 5 PiSCES changed significantly following NMDA in both tissue types (Fig 3F); for example, GluR1_PSD95 began with higher MFI in CTX, but increased in both CTX and HC; conversely Homer_PSD95 started lower in CTX, but decreased in both tissue types. For PiSCES that were significant only in CTX (Fig 3E), most were detected at low abundance in aCSF and increased with NMDA, whereas in the HC these same PiSCES were already at the higher abundance at baseline, the exception being Ube3A which remained consistently low in HC. For PiSCES that were significant only in HC (Fig 3G), most were detected at a higher abundance in aCSF and decreased with NMDA. The abundance of these PiSCES in the cortex started a lower level and trended towards a decrease with NMDA, but the magnitude of the decrease was not large enough to fulfill strict ANC∩CNA criteria. These data reveal a complex interaction between basal PIN states and activity-dependent PIN encoding of stimuli in cortex and hippocampus.

**Fig 3:**
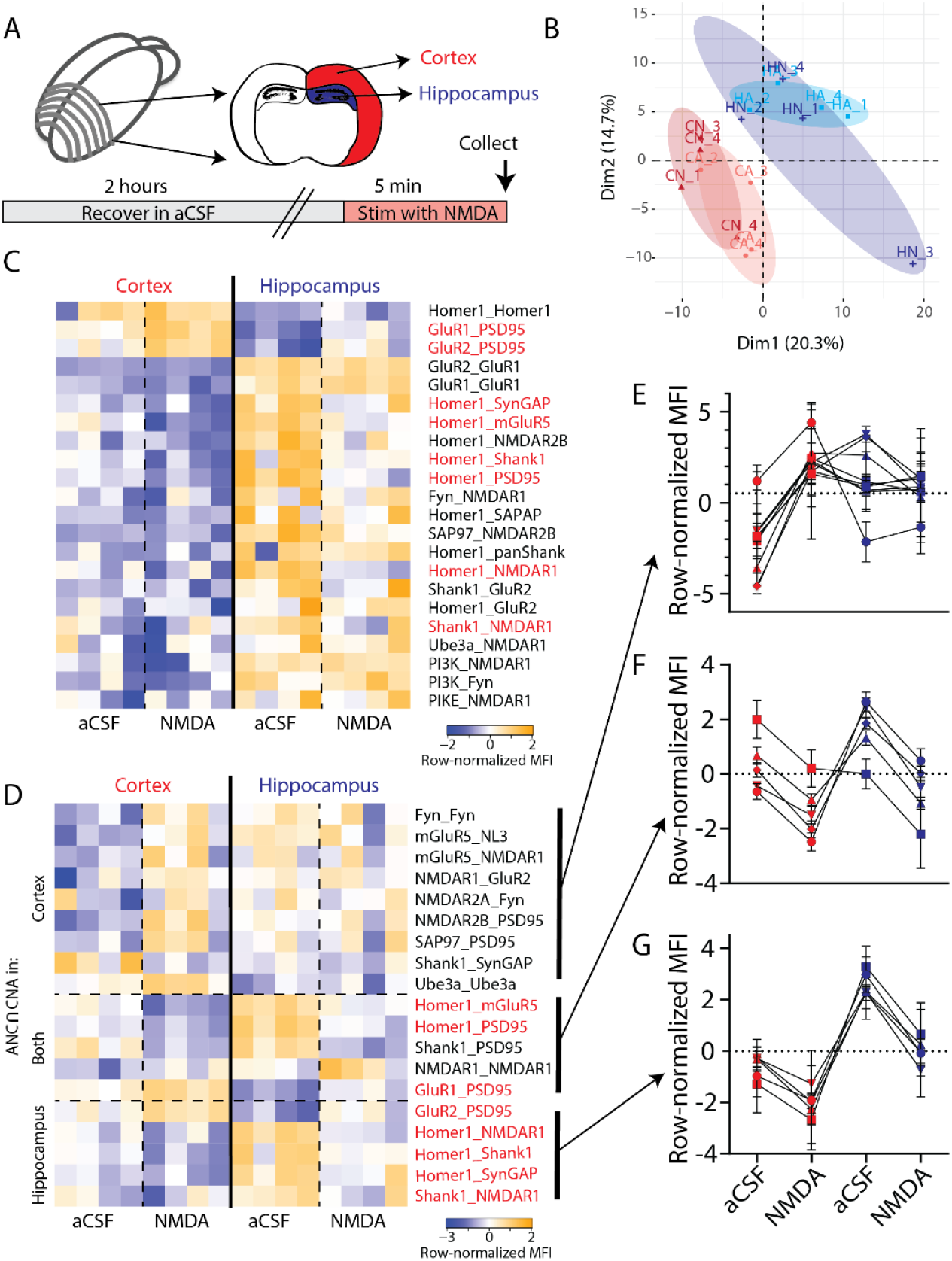
NMDA stimulation in Cortex and Hippocampus. A) Experimental design. B) PCA of hippocampal (HA and HN) and cortical (CA and CN) aCSF and NMDA groups demonstrates separation by tissue but not by stimulation. C) Heatmap of all PiSCES ANC∩CNA significant for the Cortex vs. Hippocampus comparison. D) Heatmap of all PiSCES ANC∩CNA significant for the NMDA vs. aCSF comparison. E-G) Normalized MFIs for each PiSCES ANC∩CNA significant in cortex (E), hippocampus (G), or both (F). Points represent the average normalized MFI of N=4 samples per PiSCES. Lines connect points representing the same PiSCES.

In a separate experiment, we repeated QMI after NMDA receptor stimulation in acutely-prepared brain stem slices containing the inferior olive (IO), a pre-cerebellar nucleus in the medullary brain stem. PCA showed some separation of aCSF and NMDA (Fig 4A), and ANC∩CNA identified 7 PiSCES that significantly changed with NMDA (Fig 4B), involving Homer, Shank3 and NMDARs. Batch effects inherent to QMI prevent direct comparisons of protein composition across separate experiments, so we used heatmaps that were row-normalized independently for all three tissue types to observe the behavior of PiSCES that changed significantly with NMDA in any tissue type (Fig 4C). Compared to HC and CTX, the IO was unique (Fig 4D). Zero of 19 PiSCES that were significant with NMDA stimulation in the HC or CTX changed significantly in the IO (by ANC∩CNA), even though the majority were detected. Conversely, 0/7 PiSCES that responded to NMDA in the IO responded to NMDA in HC/CTX. While many PiSCES decreased with NMDA in HC/CTX, all significantly changed PiSCES increased in the IO. Overall, these data indicate a unique PIN response to NMDA stimulation in the IO, compared to HC/CTX. Collectively, these in vitro experiments show that chemical stimulation of glutamate receptors or downstream signaling pathways causes the synaptic PIN to undergo stimulus-specific rearrangements that differ by input type and by proteomic composition of the brain area under study.

**Fig 4:**
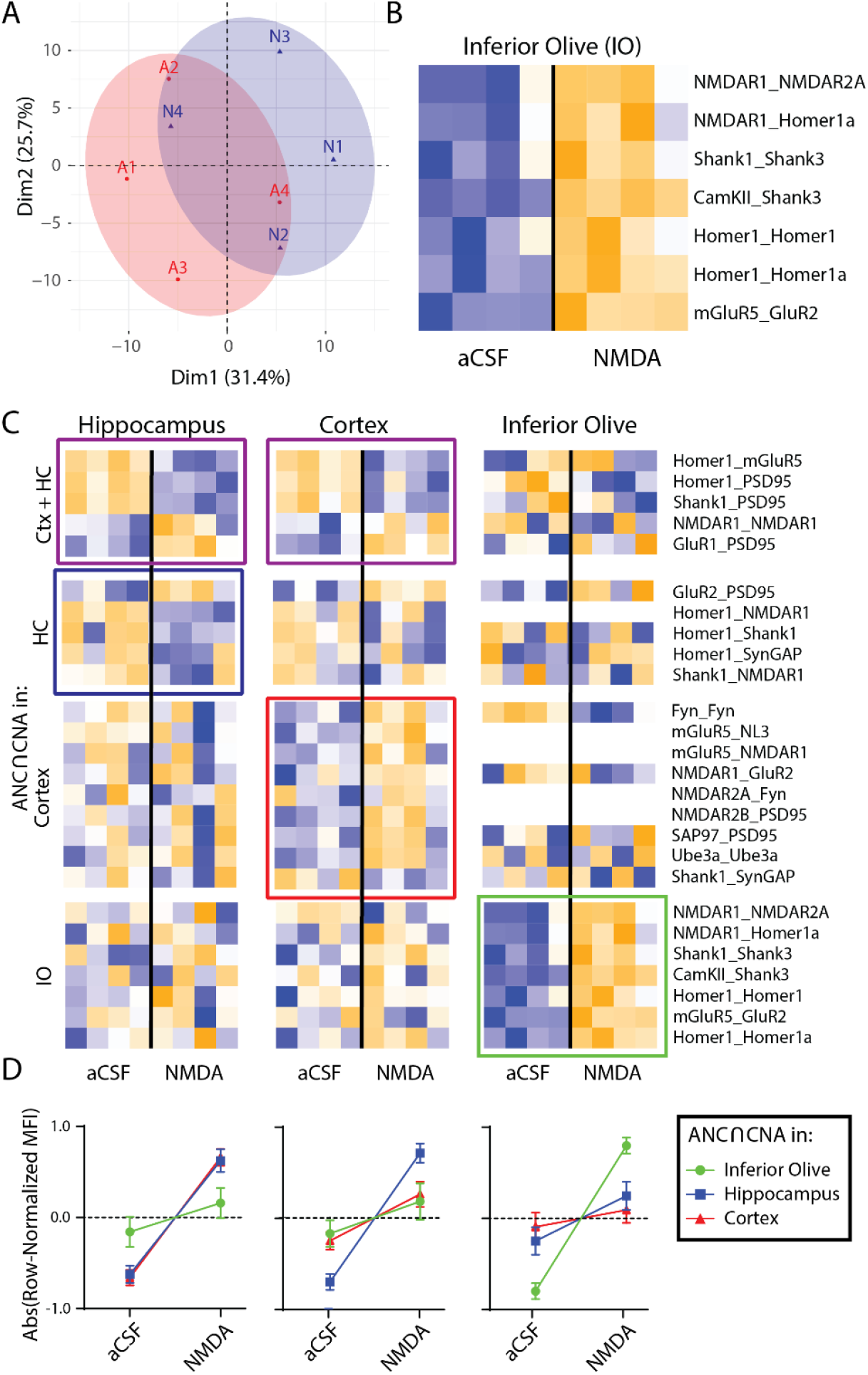
NMDA stimulation in the IO. A) PCA of aCSF (A) and NMDA (N) groups. B) Heatmap of all PiSCES ANC∩CNA significant for the NMDA vs. aCSF comparison. C) Heatmaps of all PiSCES ANC∩CNA significant for the NMDA vs. aCSF comparison in cortex, hippocampus, and IO (Cortex and Hippocampus data are the same as in Fig 3). The heatmap is row-normalized separately for each tissue to highlight overall PiSCES behavior while ignoring any baseline differences in protein expression. Boxes indicate PiSCES that are ANC∩CNA significant for that condition. D) Normalized MFIs showing the overall behavior of for all PiSCES ANC∩CNA significant in cortex, hippocampus, or IO in each tissue type.

### Different forms of associative learning elicit region- and learning-specific rearrangements in synaptic multiprotein complexes

We next asked how sensory stimuli are processed via PiSCES networks *in vivo* by inducing two forms of associative learning in awake behaving mice. These experiments employed classical eye-blink conditioning (EBC) in which an auditory conditioned stimulus (CS) preceded a periorbital electric shock unconditioned stimulus (US) that elicited a blink reflex (Fig 5A). Over repeated CS-US pairings, mice associate the tone CS with the US and acquire conditioned eye-blink responses (CRs) to the tone in anticipation of the US [45, 46]. The first paradigm was trace EBC, which is defined by a period of no stimulation (a trace interval) between CS offset and the US (Fig 5C). The second paradigm was delay EBC, which is defined by the CS extending through the CS-US interval (Fig 5D). It is well known that trace and delay EBC evoke two forms of associative learning that differ in their essential neural circuitry [47], relative involvement of the forebrain [48, 49], and dependence upon awareness (in humans [50]). The explicitly unpaired control group for behavioral and QMI comparisons consisted of mice that received a procedure in which the handling, session duration, and number of delivered CSs and US were identical to the EBC groups, except that the CS and USs were never presented together (Fig 5B). Thus, all mice experienced the same CS and US, the identical number of CS-US trials over an identical session duration, as well as identical handling and restraint. The only parameter that differed between groups was the CS-US interval.

**Figure 5:**
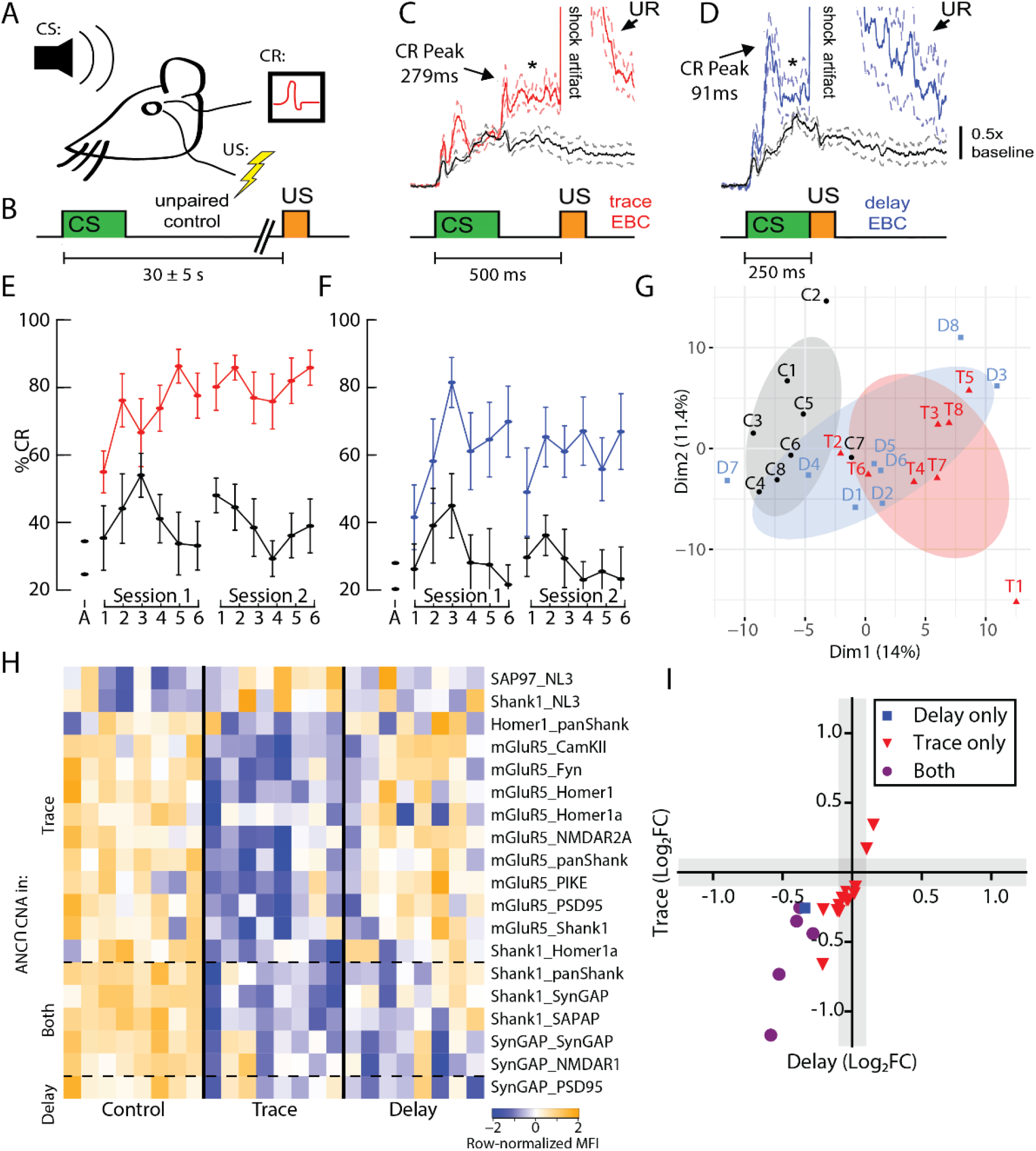
Cortex eyeblink conditioning. A) Experimental design. B) CS-US interval for the explicitly unpaired control. C,D) Average topography of CRs acquired during trace EBC (C, red) and delay EBC (D, blue) compared to responses elicited by the CS during the unpaired procedure (black). Block diagrams indicate CS-US timing for the 2 groups. Data are mean ± 1 SEM. E,F) The percent CRs exceeded the unpaired control group for both E) trace EBC (F_(1,14)_=35.8, p = 0.000) and F) delay EBC (F_(1,14)_=10.3, p = 0.006). G) PCA for the control [C], delay [D] and trace [T] conditions. H) Heatmap of row-normalized MFIs for all PiSCES ANC∩CNA significant in the delay or trace experiments.

Mice receiving trace (Fig 5E) or delay (Fig 5F) EBC showed robust CR acquisition over 2 EBC sessions, as evidenced by a percentage CRs that greatly exceeded the CS responding of mice that received the explicitly unpaired control procedure. The use of an explicit tly unpaired control was important to confirm that CR acquisition was specifically due to associative learning and was not due to non-associative influences such as sensitization, an increase in baseline responding, or pseudoconditioning. Figure 5C and D demonstrate the average topography of CRs acquired to the CS under the trace and delay EBC paradigms, indicating that their temporal kinetics adapted to the CS-US interval; responses were significantly larger than responses elicited unconditionally by the tone CS in the unpaired group, both of which are hallmarks of associative learning during EBC. Immediately following the second training session, the mice were rapidly anesthetized with isoflurane, decapitated, and the medial prefrontal cortex (mPFC), HC and IO were dissected to allow activity-dependent PIN rearrangements to be quantified by QMI.

In the mPFC, PCA showed separation of Trace eyeblink EBC from unpaired controls across PC1, while Delay EBC was intermediate (Fig 5G). ANC⋂CNA identified 19 PiSCES that were significantly different between the trace or delay EBC and controls; 18 that were significant for trace EBC, 6 for delay EBC, and 5 that were common to both learning paradigms (Fig 5H). Similar to NMDA stimulation experiments, dissociations among Shank- and SynGAP-containing PiSCES were prevalent. Interestingly, several mGluR5-containing PiSCES decreased strongly in trace, but did not change in delay EBC. When we directly compared activity-dependent changes using an X-Y plot (Fig 5I), we observed a strong correlation (R^2^= 0.71) between the two conditions, with the slope of the regression line (1.32) confirming that PiSCES activation was stronger in trace EBC. Overall, these data demonstrate that stimulation of the mPFC by a learning paradigm induces dissociation of PiSCES, especially those containing Shank, SynGAP and mGluR5. The two learning paradigms, which differed only in the timing of the CS-US, activated an overlapping set of PiSCES, but the cortex-dependent paradigm produced stronger and more extensive PiSCES activation.

In the HC, PCA did not clearly distinguish the groups (Fig 6A). However, 4 PiSCES were ANC∩CNA significant for trace EBC and 22 were significant for delay EBC (Fig 6B). Shank3- and NMDAR-containing PiSCES were most prominently activated for delay EBC, with clear increases in co-associations among NMDARs, FYN, NL3, and the Homer-Shank scaffold demonstrating strong upregulation of NMDAR-containing complexes. For Trace EBC, we observed weak and variable mGluR5_Homer1 and SynGAP_PSD95 dissociation, as well as changes in PI3K and Fyn self-association. When we compared activity-dependent changes using an X-Y plot (Fig 6C), the majority of PiSCES that were significant for one form of learning fell below the limit of detection for the other form of learning, although the most strongly activated PiSCES in delay EBC showed a trend toward increasing in trace EBC.

**Fig 6:**
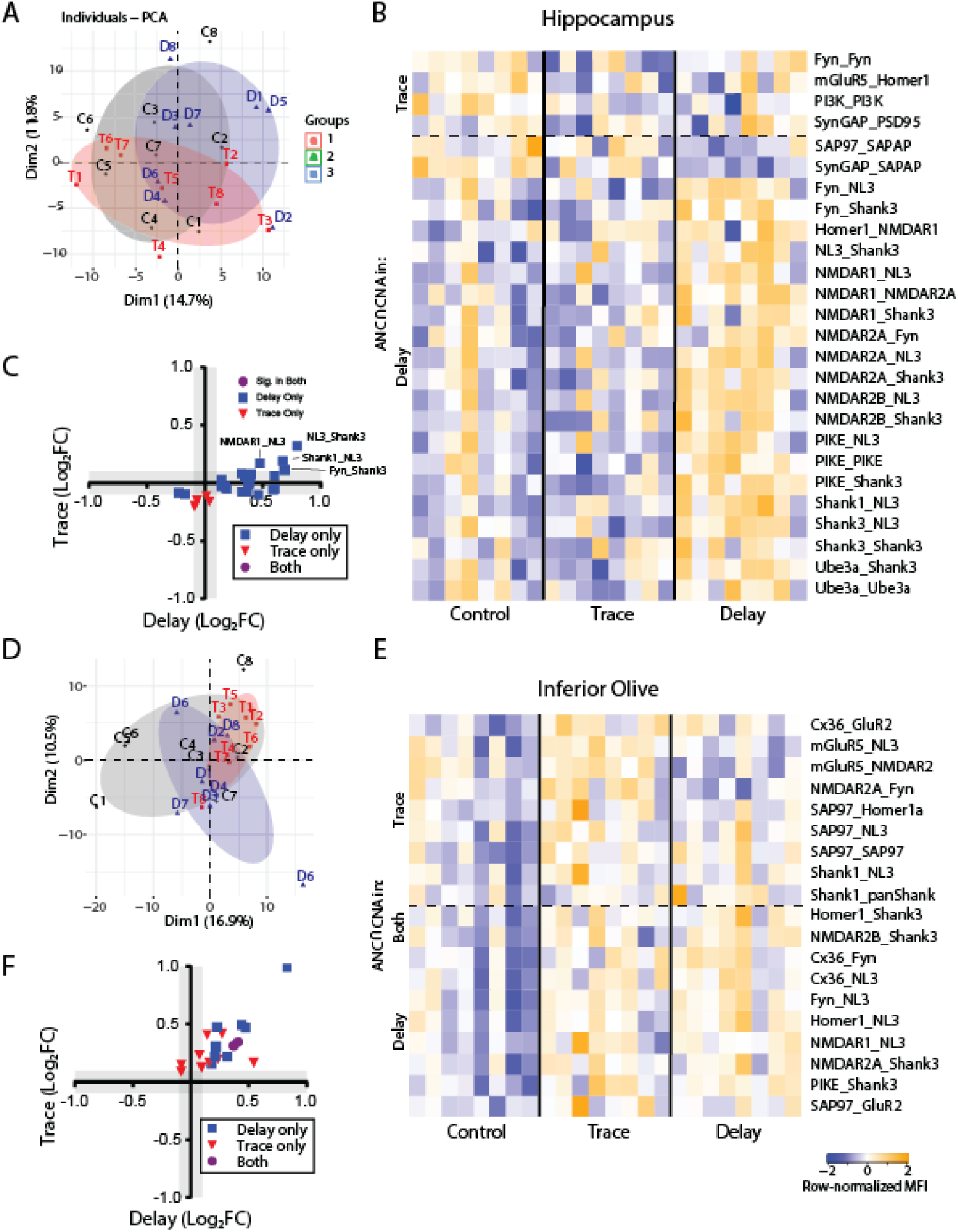
Hippocampus and IO eyeblink conditioning. A) PCA for the control [C], delay [D] and trace [T] conditions in Hippocampus. B) Heatmap of row-normalized MFIs for all PiSCES ANC∩CNA significant in Hippocampus for the delay or trace experiments. Data are expressed as row-normalized MFI. C) X-Y plot comparing the log_2_ fold change of all PiSCES that were ANC∩CNA significant for either Delay (blue), the Trace (red) or both experiments (purple) in Hippocampus. D) PCA for the control [C], delay [D] and trace [T] conditions in the IO. E) Heatmap of row-normalized MFIs for all PiSCES ANC∩CNA significant in the delay or trace experiments in the IO. Data are expressed as row-normalized MFI. F) X-Y plot comparing the log_2_ fold change of all PiSCES that were ANC∩CNA significant for either Delay (blue), the Trace (red) or both experiments (purple) in the IO.

We next analyzed the IO, a pre-cerebellar nucleus in the medulla oblongata that is required for both trace and delay EBC [51, 52]. As for HC, PCA showed no clear separation between control and stimulated conditions (Fig 6D), but ANC⋂CNA identified 8 and 10 interactions that were significantly different following delay and trace EBC, respectively, with two PiSCES significant in both conditions (Fig 6E). All significant PiSCES involved increases in the magnitude of interactions, and the proteins involved - e.g. Shank3, NL3, and SAP97 - were different from those observed in mPFC and HC experiments. When we compared activity-dependent PiSCES between trace and delay EBC in the IO (Fig 6F), we found a moderate correlation (R^2^ = 0.56), suggesting a common set of qualitatively similar changes in PiSCES, with the slope of the regression line (0.71) indicating stronger activation following delay EBC. Overall, these data reveal that trace and delay EBC elicit qualitatively similar within-region changes in the IO and mPFC, while in HC, a different protein-state-dependent encoding paradigm differentiates trace from delay EBC. Moreover, these data show how two distinct forms of associative learning elicit brain-region-specific PIN rearrangements *in vivo*.

## Discussion

Far from a static ‘scaffold’, the post-synaptic density is a dynamic, liquid-liquid-phase-separated [53–56] structure that rapidly responds to incoming synaptic activity by altering its molecular composition via regulation of protein-protein interactions. Certainly, individual interactions, such as recruitment of AMPARs to PSD95 scaffolds [57, 58] or dissociation of mGluR from Homer EVH1 domains [33, 34] are well-characterized, but the molecular logic that allows a change in protein interactions to translate into functional cellular ‘calculations’ (Fig 1A) remains understudied, partly due to technical limitations. Here, using QMI, we investigated how networks of interacting proteins encode incoming synaptic stimuli, and how that encoding differs between brain regions.

Cells can use diverse mechanisms to encode signaling inputs, and to perform calculations that produce an appropriate response to environmental conditions. The modular hypothesis, first developed by Pawson and colleagues, suggested that different combinations of proteins recruited to signaling complexes could define different signals [36, 59], but unambiguous examples of this intuitive style of signaling have been experimentally elusive. Differences in the temporal dynamics of signals have also been proposed to mediate cellular decisions; notably “weak and fast” vs. “strong and sustained” intracellular Ca2+ transients have been proposed to mediate the LTD vs. LTP decision [60], in a mechanism that may involve auto-inhibition of CamKII [26, 61]. In an elegant series of optogenetic experiments, Toettcher and colleagues demonstrated that the ERK signal transduction cascade acts as a temporal filter, detecting sustained activation of ERK signaling while rejecting more transient ERK phosphorylation [40]. In addition to qualitative and kinetic logic, some systems simply use the magnitude of signalosome activation to encode different stimuli. In T cell development, the magnitude of activation of a single PIN downstream of the T cell receptor determines whether a cell chooses one of two diametrically opposed outcomes-continued development or apoptotic cell death [39]. Finally, computational modeling of BMP ligand/receptor interactions revealed a complex and unintuitive combinatorial logic system, in which peak receptor activation is achieved at specific concentrations of ligands determined by receptor sub-type, and different combinations of receptors and ligands allow a diversity of cellular logic responses based on concentration gradients [62].

In the brain, the molecular logic rules that allow neurons to distinguish between signaling inputs and synthesize coordinated cellular responses are largely unknown. Here, we first used a simple cell culture model to explore the encoding of two major glutamate receptor classes, metabotropic vs. NMDA receptors. We found that qualitative, quantitative and kinetic mechanisms combine to differentiate the signals. As quickly as 30s, a single module was activated uniformly downstream of both receptors. This is congruent with Ca2+-dependent signaling, as intracellular Ca2+ is induced by both NMDA (via direct conduction) and DHPG (via IP3R and release from the ER) [63, 64]. At 5 and 15 min, this same module activated more intensely in NMDA and glutamate conditions, but quickly returned to baseline in DHPG, demonstrating kinetic differentiation of signals consistent with a ‘weak-and-fast’ vs. ‘strong-and-sustained’ Ca^2+^ flux model of LTD vs. LTP [60]. In stark contrast, cLTP did not resemble either NMDA or DHPG application, and instead produced a distinct pattern of PiSCES changes, which were often in the opposite direction compared to NMDA or glutamate. While bath application of NMDA has been reported to cause an LTD-like response [65, 66], AMPA receptor-scaffold co-associations increased with NMDA/glutamate (implying an increase in synaptic strength) but did not change with cLTP. A limitation of our study is that we did not measure the electrophysiological correlates of our treatments, so we cannot conclude if receptor activation induced an LTP- or LTD-like response. We can conclude that, in a cultured neuron system, the synaptic PIN can obtain multiple independent states (at least three) that correspond to unique input types. These states are defined by differences in composition, intensity and kinetics of PIN activation.

### Proteome-dependent PIN activation

The core proteins targeted by our QMI panel co-exist in multiprotein complexes throughout the brain. However, the proteome of neurons varies by brain area. Accessory proteins in complex with our QMI targets, or splice variants of the targets themselves, may allow different brain regions to perform unique molecular computations, suited for the particular circuitry of the area. To explore this idea, we directly compared an identical stimulus in two brain regions with known differences in proteome composition - the HC and CTX. When NMDA was applied, the PIN response was broadly similar, involving decreased Homer-Shank scaffold interactions and increased AMPA and PSD95 PiSCES. However, differences in the levels of these PiSCES at baseline between the two areas meant that the dynamic range available for a given PiSCES to change differed. Conversely, the response to NMDA in the IO was unique. PiSCES involving Shank3 and NMDARs were strongly upregulated, as well as a PiSCES containing mGluR5 and GluR2, which may be linked by Shank3 scaffolds [67]. In HC/CTX, these PiSCES did not change with NMDA. The different ratios of Shank isoforms expressed in these three tissues may partially explain these differences, since Shank3 is highly expressed in the IO [68]. Overall, these data demonstrate that baseline proteomic differences allow glutamate receptors and scaffolds to modify their interactions in response to NMDA stimulation in very different ways, allowing for a diversity of computations across brain regions.

### PIN activation underlying associative learning

The EBC paradigms allowed us to present mice with three distinct types of information, while using identical sensory stimuli. Every mouse received the same number of CS and US presentations, but their timing determined which type of conditioning the animal received. Remarkably, this sole difference in timing resulted in different PIN responses within the same brain areas. In the mPFC, the PiSCES activated by EBC were similar to those activated by NMDA/glutamate in neocortical neurons in culture (dissociation of mGluR5, Homer, Shank and SynGAP), with the intensity of activation and the involvement of mGluR5 distinguishing delay vs. trace EBC. Consistent with this result, inhibition of NMDA receptors the medial prefrontal cortex has been shown to severely inhibit classical EBC [69, 70]. By comparison, in the HC the PiSCES activated in trace vs. delay EBC were non-overlapping. Trace conditioning weakly activated Fyn, PI3K, mGluR5_Homer1 and SynGAP_PSD95, PiSCES associated with in vitro NMDA/glutamate stimulation. We speculate that these data are consistent with activation of a sparse population of cells in a background of inactive cells, as might be expected for a hippocampal-dependent task, although other explanations certainly exist. However, delay conditioning, which is not dependent on the HC, elicited a strong PIN response consisting of up-regulation of NMDAR, Shank, Fyn, and NL3 complexes. This pattern of PIN activation was not observed in our prior agonist-stimulation experiments, nor have we observed a similar pattern in response to homeostatic scaling, in vitro or in vivo [71]. Finally, the IO, which is essential for the performance of CRs in both trace and delay EBC [72, 73], also activated a PIN consisting of Shank3, Fyn, and NL3 PiSCES in response to both EBC types. Overall, these results demonstrate a remarkable diversity of PIN responses to natural stimuli, which may serve as a basis for the diversity of functions that neuronal subtypes can perform across the brain.

Why might the HC have responded so strongly to delay EBC, when it is required for trace, but not delay, EBC? Perhaps the hippocampal activation observed here serves an alternative purpose, outside the acquisition of the CR. Rats with hippocampal lesions only exhibit deficits in a conditional EBC paradigm when they must learn to blink only if the CS is preceded by another stimulus [74]. Humans with hippocampal-temporal lobe anterograde amnesia can acquire a CR, but are unable to describe it [75], and patients with hippocampal-medial temporal lesions acquire a CR, but have impaired conditional discrimination in EBC [76, 77]. While PIN activation in the HC observed here may not reflect acquisition of the CR, perhaps it is an underlying molecular event in hippocampal-dependent declarative memory.

### Dysfunction of PIN dynamics in neurological disorders

Synaptic dysfunction is causally associated with numerous neurological disorders including Autism Spectrum disorders (ASDs), schizophrenia, and bipolar disease [78–82]. The synaptic QMI panel itself was designed to target a network of autism-linked gene products [37] in order to understand how mutations in any one member of a highly interconnected PIN produces a similar human phenotype in the form of ASD [71]. Indeed, we and others have previously demonstrated that mice carrying autism-linked mutations in Shank3 or Homer1 show similar disruptions in specific types of plasticity, such as homeostatic scaling [71]. As our understanding of ASD biology improves, many (although not all) ASD-linked genes can be placed in a pathway that translates synaptic activity into coordinated changes in neuronal protein networks, over developmental time [82]. The work presented here further demonstrates that ASD-linked proteins including Homer, Shank, SynGAP, NMDAR2B and mGluR5 are intimately involved in neuronal computations downstream of both chemical and sensory inputs.

It is hoped that as we better understand the mechanisms of pathogenic mutations, we will develop treatments to correct molecular deficits and improve patient outcomes. However, our data highlight a significant challenge in developing a therapeutic: how does one correct signaling deficits that are not consistent across the entire brain? Region specific electrophysiological deficits in numerous models of ASD are well established (e.g. [83]). Here, we demonstrate that even the rules governing basic signal transduction networks differ between brain regions when an animal is performing a simple task. That complexity is likely to be greatly amplified when the animal performs two or more different tasks. Thus, correcting a signaling deficit in one brain region could have a confounding or even contradictory effect on other regions or other behaviors. While this may complicate therapeutic development, recognizing this diversity may be critical to our integrative understanding of brain function in health and disease.

## Acknowledgments

The authors would like to thank Dr. Adam Schrum, and members of the Smith lab, especially Dr. Whitney Heavner and Devin Wehle for helpful discussions concerning the data and manuscript. The authors also thank Liza Severs and the Ramirez lab for their gift of Ai95D and Vglut2-ires-cre mice.

## Author contributions

JL performed all experiments with technical assistance from EG and ZZ, except for the eyeblink conditioning experiments performed by ZZ and JW, and the two-photon imaging performed by KT. JL, KT, SEPS, and JPW analyzed the data. JL, SEPS and JPW wrote the manuscript. All authors read and approved the manuscript.

## Materials and Methods

### Cell culture and Neuronal Stimulation

Primary cultures of cortical neurons were prepared as previously described [29]. Briefly, whole cortex from P0-P1 mouse neonates was dissociated using papain (Worthington) and plated at a density of 1.0×10^6^ cells/mL onto 6-well plates treated with poly-D-lysine. Cells were cultured in Neurobasal medium supplemented with 2% B27 and 0.5mM GlutaMax (ThermoFisher) and kept at 37°C, 5% CO2 for 18-21 days. After 3-5 DIV, 5-fluoro-2’-deoxyuridine was added to a final concentration of 5 μM to inhibit glial proliferation.

For direct activation of glutamate receptors, neurons were treated with Glutamate (100 μM), NMDA/glycine (100/1 μM), DHPG (100 μM) or control HEPES-aCSF (129 mM NaCl/ 5 mM KCl/ 2 mM CaCl2/ 30 mM Glucose/ 25 mM HEPES; pH 7.4) for up to 5 minutes. For longer time periods, aCSF was removed and replaced with conditioned media. Glutamate (Sigma, G1626; 100 μM or 1 μM) was prepared in HEPES-aCSF. NMDAR agonists NMDA (Sigma, M3262; 100 μM) + glycine (Fisher Scientific, Pittsburgh, PA, USA, BP-381; 10 μM) were dissolved in magnesium-free HEPES-aCSF in the presence of non-NMDA glutamate receptor blockers (CNQX, Tocris, 0190, 40 μM; Nimodipine, Tocris, 0600, 1 μM; LY-367385, Sigma, L4420, 100 μM; MPEP, Tocris, 1212, 10 μM). Type I mGluR agonist DHPG (Tocris, 0805; 100 μM) was dissolved in HEPES-aCSF in the presence of non-mGluR glutamate receptor blockers (CNQX, Tocris, 0190, 40 μM; Nimodipine, Tocris, 0600, 1 μM; D(-)-2-Amino-5 phosphonopentanoic acid (APV), Sigma, A5282, 50 μM). For chemical induction of LTP, neurons were treated as previously described [44]. Briefly, neurons were treated with cLTP solution containing Forskolin (Tocris, 1099, 50 μM), Rolipram (Tocris, 0905, 0.1 μM), and Picrotoxin (Tocris, 1128, 50 μM) dissolved in HEPES-aCSF or control HEPES-aCSF for 30 minutes. Following treatment, cells were immediately placed on ice and rapidly harvested by scraping in lysis buffer [150 mM NaCl, 50 mM Tris (pH 7.4), 1% NP-40, 10 mM sodium fluoride (Sigma, 201154), 2 mM sodium orthovanadate (Sigma, 450243), Protease/phosphatase inhibitor cocktails (Sigma, P8340/P5726), followed by incubation in lysis buffer for 15 min. Lysate was centrifuged for 15 min at 15,000G to remove nuclei and debris, and protein concentration in the supernatant was determined using a Pierce BCA kit (Pierce, Rockford, IL, USA, 23225).

### Animals

CD-1, Vglut2-ires-cre (stock 01963), and Ai95(RCL-GCaMP6f)-D or Ai95D (stock 028865) mice were originally obtained from The Jackson Laboratory (Bar Harbor, Maine) and maintained in an in-house breeding colony. All mice were separated by sex, and housed with littermates, with no more than five mice/cage. Food and water was provided ad libitum. For slice experiments, only p21-30 mice (both male and female) were used. The use and care of animals was approved by the Office of Animal Care at the Seattle Children’s Research institute (protocol# IACUC0072) and complied with all relevant guidelines and regulations.

### Slice preparation and treatment

Mice were deeply anesthetized with Isofluorane, brains were removed, and coronal cortical and hippocampal slices were sectioned at 400 μm thickness using a vibratome. Slices were immediately hemisected with a sharp razor blade and each half placed in an alternate treatment group with treatment groups being arbitrarily assigned. For IO slices, brains were removed, hemisected coronally at the midbrain and the cerebellum was removed, and slices sectioned at 400 μm thickness using a vibratome. The IO was then microdissected away from surrounding tissue. Slices were initially recovered in NMDG protective recovery solution (93 mM NMDG, 2.5 mM KCl, 1.2 mM NaH2PO4, 30 mM NaHCO3, 20 mM HEPES, 25 mM glucose, 2mM thiourea, 5mM Na-ascorbate, 3mM Na-pyruvate, 0.5 mM CaCl2.4H2O, and 10 mM MgSO4.7H2O; titrated to pH 7.4 with concentrated hydrochloric acid) for 10–15 min at 32–34°C, then transferred to a modified HEPES holding solution [92 mM NaCl, 2.5 mM KCl, 1.2 mM NaH2PO4, 30 mM NaHCO3, 20 mM HEPES, 25mM glucose, 2mM thiourea, 5mM Na-ascorbate, 3mM Na-pyruvate, 2mM CaCl2.4H2O, and 2mM MgSO4.7H2O; pH 7.4] for an additional 60–90 min recovery at room temperature. Slices were then incubated in HEPES-aCSF x 1h at 32–34°C to equilibrate, and subsequently stimulated with NMDA/glycine (100/1 μM) or HEPES-aCSF control, similar to cell culture experiments. Following stimulation, slices were homogenized in ice cold 1% NP-40 lysis buffer using 12 strokes of a glass-teflon homogenizer.

### Eyeblink conditioning

EBC was performed as described in [45]. Adult mice of both sexes were implanted with stimulation and recording electrodes under Isoflourane anesthesia and were allowed to recover for >1 week before experimental sessions commenced. Head-restrained, freely-walking mice (N = 24) were then acclimated to the conditional apparatus and conditioned over two days. The CS was a 2-kHz, 85-dB, 250-ms, free-field tone. The US was an 800-μA, biphasic 100-ms shock to the right periorbital region. Eyeblinks were recorded by intramuscular EMG obtained from the right superior orbicularis oculi muscle (bipolar electrode, 1-kHz sampling, baseline rectified, 200-Hz low-pass filtered). A CR was defined as an EMG signal within the CS-US interval exceeding 6 standard deviations of a stable, 100-ms pre-CS baseline with an onset latency greater than 35 ms and duration exceeding 15 ms. Comparisons of CR acquisition in trace and delay EBC groups were made to the same unpaired control group using an identical “CS-US” interval as the conditioning group. Mice receiving trace or delay EBC experienced 60 CS-US conditioning trials per hour-long session, each trial consisting of a 250 ms CS and a 100 ms US. Mice receiving trace EBC experienced a 500 ms CS-US interval (250 ms trace interval) while mice receiving delay EBC experienced a 250 ms CS-US interval (no trace interval). CSs and USs during the explicitly unpaired paradigm were delivered alone (no more than 3 consecutive trials of the same type) at half the EBC intertrial interval (30 +/- 5 s). Immediately following the second training session, the mice were rapidly anesthetized with isoflurane, decapitated, and the medial prefrontal cortex (mPFC), HC and IO were dissected to allow activity-dependent PIN rearrangements to be quantified by QMI.

### Quantitative Multiplex co-immunoprecipitation (QMI)

QMI was performed as described previously [29, 38, 41]. Briefly, a master mix containing equal numbers of each antibody-coupled Luminex bead was prepared and distributed to lysates containing equal amounts of protein and incubated overnight on a rotator at 4°C. The next day, beads from each sample were washed twice in cold Fly-P buffer (50mM tris pH7.4, 100mM NaCl, 1% bovine serum albumin, and 0.02% sodium azide) and distributed into twice as many wells of a 96-well plate as there were probe antibodies (for technical replicates). Biotinylated detection (probe) antibodies were added to the appropriate wells and incubated at 4°C with gentle agitation for 1 hour. The resulting bead-probe complexes were washed 3 times with Fly-P buffer, incubated for 30 minutes with streptavidin PE on ice, washed another 3 times, resuspended in 125μl ice cold Fly-P buffer, and processed for fluorescence using a customized refrigerated Bio-Plex 200 [41]. Antibody clone names catalog numbers are the same as in [29].

### Data Analysis

CNA: Modules of PiSCES that co-varied with experimental conditions were identified using CNA as described in [29, 41]. Briefly, bead distributions used in ANC were collapsed into a single MFI for every PiSCES and averaged across technical replicates for input into the WGCNA package for R [84]. PiSCES with MFI < 100 were removed, and batch effects were corrected using COMBAT [85]. For timecourse analysis, different timepoints were further batch-corrected using COmbat CO-Normalization Using ConTrols (COCONUT) [86]. Power values giving the approximation of scale-free topology were determined using soft thresholding with a power adjacency function. The minimum module size was always set to between 10 and 12, and modules whose eigenvectors significantly correlated with an experimental trait (p<0.05) were considered “of interest.” PiSCES belonging to module of interest and whose probability of module membership in that module was < 0.05 were considered significantly correlated with that trait.

PCA: PCA was performed on Post-COMBAT, log2 transformed MFI values in R studio using the prcomp function.

ANC: High-confidence, statistically significant differences in bead distributions between conditions for individual PiSCES, after correcting for multiple comparisons, were identified using ANC as described in [41].

ANC∩CNA: PiSCES that were significant by both ANC and CNA for a given experimental condition were considered significantly altered in that condition.

### TUNEL assay and immunocytochemistry

For TUNEL staining and IHC, cortical cells were cultured as described above with the following exceptions: cells were plated at 1.0 x 10^6^ cells/mL on glass coverslips treated with poly-D-lysine in 24 well plates. Detection and quantification of apoptosis in cell cultures was determined using a *In Situ* Cell Death Detection Kit (Roche, 11684795910) according to the manufacturer’s instructions. Briefly, cells were fixed with 4% paraformaldehyde x 1 hour at room temperature, rinsed with PBS, and incubated in permeabilization buffer (0.1% Triton X-100 in 0.1% sodium citrate freshly prepared). Cells were then washed 3x with PBS and stained with TUNEL reaction mixture x 60 min at 37°C in the dark. For negative controls, fixed cells were incubated in Label solution instead of the TUNEL mixture. For positive controls, permeabilized cells were incubated with DNAse I recombinase (3-3000 U/ml in 50 mM Tris-HCL, pH 7.5, 1/ mg/ml BSA: Worthington, LK003170) prior to incubation with TUNEL mixture.

Following TUNEL staining, cells were treated with antibodies towards β3-tubulin (1:500, Biolegend, 801201) and NeuN (1:1000, Abcam, ab177847) overnight at 4°C. The following day, cells were washed 3x with PBS and incubated Alexa Fluor® 594 AffiniPure Goat Anti-Rabbit IgG (1:10,000, Jackson ImmunoResearch, 111-585-003) and Alexa Fluor® 647 AffiniPure Goat Anti-Mouse IgG (1:10,000, Jackson ImmunoResearch, 115-605-003) x 60 min at RT. Cells were then washed 3x with PBS and mounted in Fluoromount G Mounting Medium, with DAPI (Invitrogen, 00495952). 4 pictures of each coverslip were taken at random, and the NeuN positive cells that were also TUNEL positive was quantified in ImageJ, with a minimum of 500 cells/slip being counted.

### 2-photon Ca^+^ imaging

For Ca^+^ imaging experiments, primary cortical neurons were cultured on glass coverslips from P0 pups born from crossing Ai95D and Vglut2-ires-cre mice (as described above). The resulting neurons expressed the calcium indicator GCamp6F in Vglut2+ neurons. Two-photon Ca^+^ imaging was performed using an Olympus FV1000MPE multiphoton laser scanning microscope equipped with a Mai Tai Deepsee Laser. All experiments were performed using a 25x objective immersed in aCSF solution and a 2x digital zoom. For imaging, glass coverslips were placed in a recording chamber, and perfused with oxygenated aCSF at a flow rate of 5-6 ml/min. Acquisition protocols consisted of ~35-minute time-lapse sequences measuring changes in the levels of fluorescence using a GFP filter (wavelengths 457-538). To quantify changes in fluorescence intensity, the regions of interest (including both cell body and processes) were selected and mean pixel intensity was measured using the Flowview program. The data was plotted as ΔF/baseline (defined as average fluorescence in the minute preceding stimulation) relative to time. For stimulation, glass coverslips were perfused with aCSF containing either glutamate (100 μm), NMDA/glycine (100/1 μm), or DHPG (100 μm) for 5 minutes, after which coverslips were perfused with fresh aCSF lacking agonist. At 30 minutes, coverslips were perfused with high K^+^ aCSF (55 μm). Statistical significance was determined using a Student’s *t-test*.

## Supplementary Material

**Supplementary Figure 1:**
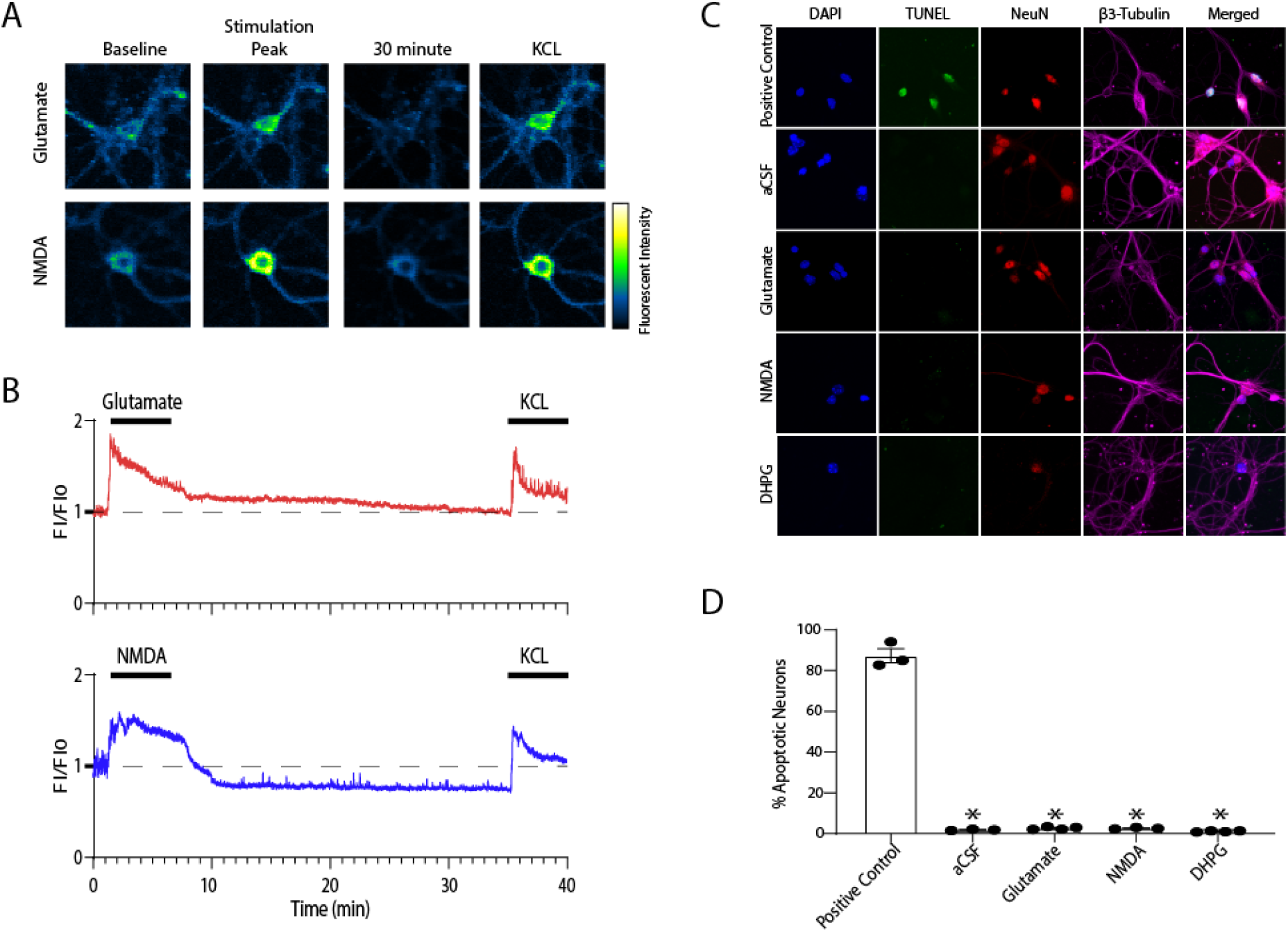
NMDA and DHPG treatments do not induce excitotoxicity. A) Example of Ca2+ imaging experiment showing baseline, stimulation peak (~2-3 minutes following NMDA), 30 minutes after wash-out, and the peak of a second KCL stimulation. B) Averaged FI/F0 traces for glutamate and NMDA stimulations. N= 28-29 neurons. C) TUNEL staining showing lack of apoptosis two hours following aCSF, glutamate NMDA or DHPG treatment. D) Quantification of data shown in C, N=3-4 experiments per condition.

**Supplementary Figure 2:**
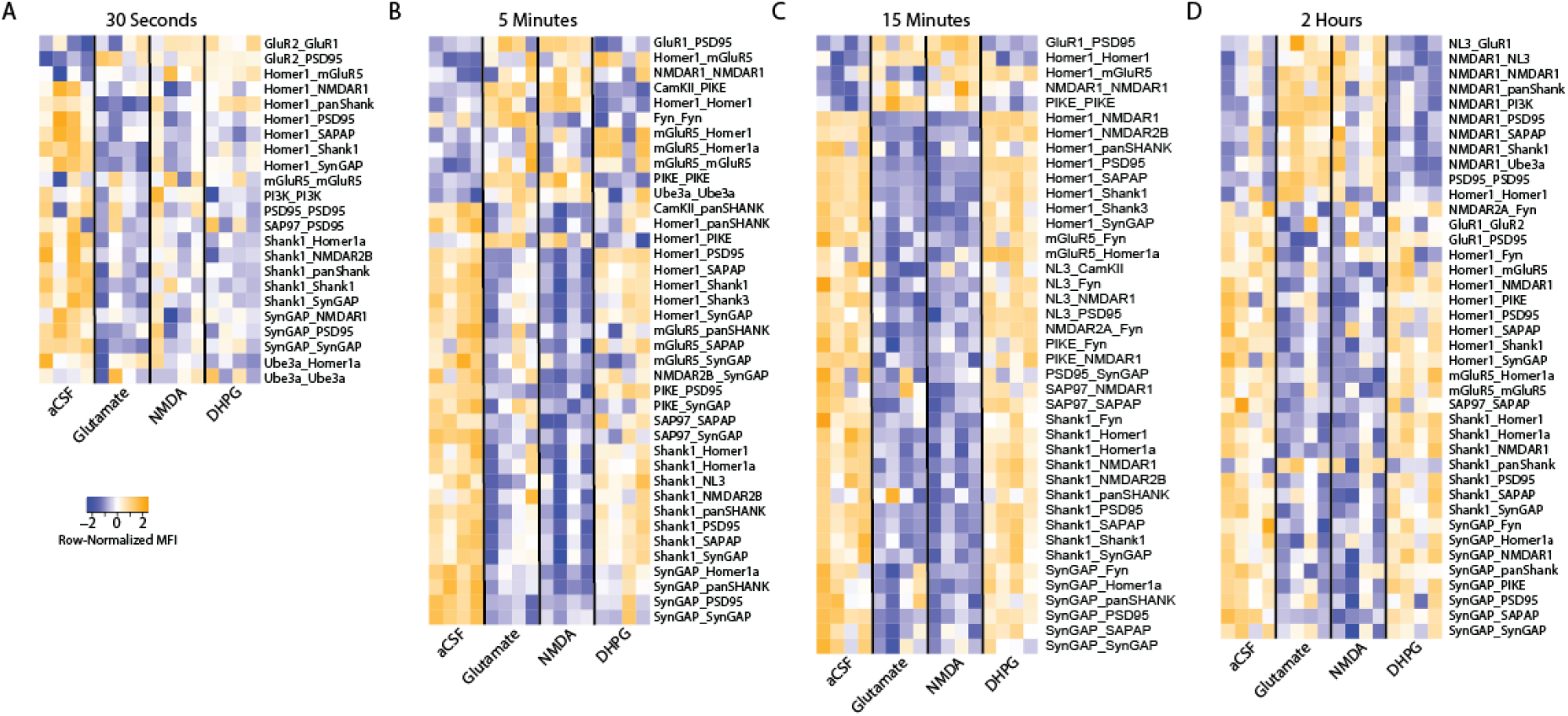
Heatmap of row-normalized MFIs for all PiSCES ANC∩CNA significant at each of the timepoints shown in **Figure 1.** Data are expressed as row-normalized MFI.

